# Predation of small nocturnal primates (*Microcebus*) on Madagascar; insights into predator preferences in Andohahela National Park

**DOI:** 10.64898/2026.08.22.746414

**Authors:** Sam Hyde Roberts, J. Carolina Segami, Vatosoa J. N. Harinala, Balsama Rajemison, Evatiana Rasoarinoro, Steven M. Goodman, Anne D. Yoder

## Abstract

Predation is a major selective force shaping lemur behaviour and population dynamics, yet direct observations remain rare, particularly for small nocturnal species. Here, we document predation on mouse lemurs (*Microcebus* spp., Cheirogaleidae) in Andohahela National Park, south-eastern Madagascar, and using complementary evidence from radio-telemetry and owl pellet analyses. Two radio-collared individuals were confirmed as prey of endemic snakes, *Ithycyphus oursi* and *Madagascarophis meridionalis* (both Pseudoxyrhophiidae), representing the first documented records of *Microcebus* predation by these species. Examination of owl pellets and prey remains from three sites and from three owl species revealed a single predation event by *Asio madagascariensis* (Strigidae), likely involving *M*. *tanosi*, no evidence of mouse lemur predation by *Tyto alba* (Tytonidae) despite high local prey availability, and a single and first predation record from *Athene superciliaris* (Strigidae). Material from *A. superciliaris* roosts was otherwise dominated by invertebrates, indicating that primate predation is likely opportunistic. Together, these findings expand the known predator guild of mouse lemurs and suggest that snake predation is rarely detected, and its contribution to mouse lemur mortality and population dynamics is likely underestimated. Our findings demonstrate how combining behavioural field observations with dietary evidence can uncover otherwise undetected predation events and clarify predator prey relationships in nocturnal primates.

## Introduction

Evidence accumulated over recent decades, indicates that predation has played a central role in shaping lemur behavioural ecology and population dynamics (Goodman et al., 1993; Karpanty, 2003; Meador et al., 2019; Goodman and Ganzhorn, 2022). Since the vast majority of lemur species experience some degree of predatory pressure (Goodman and Ganzhorn, 2022) and play important ecological roles as seed dispersers, pollinators, and foragers within Malagasy ecosystems (Sussman and Raven, 1978; Sato, 2012; Eppley et al., 2022; Ramananjato, 2025), understanding how predation influences their population dynamics is of particular interest. Although the full extent of predator-prey interactions remains incompletely understood, predators from across the major Malagasy predator groups are known to prey on the smallest lemurs, the mouse lemurs (Cheirogaleidae: *Microcebus*). With 19 recognized species (Van Elst et al., 2025), weighing between 30–90 g (Mittermeier et al., 2023) and distributed widely across Madagascar, mouse lemurs are exposed to a wide and varied assemblage of predators. Since predation events are seldom observed directly, much of the available evidence derives from indirect sources, including predator remains (e.g., pellets or scats) and opportunistic field observations. Nevertheless, the inventory of known *Microcebus* predators continues to expand with increasing field research (Ramananjato et al., 2025). Presently, predation records exist for 11 *Microcebus* species (Table 1), with some 15 predatory species documented (Fichtel, 2016; Goodman and Ganzhorn, 2022). These findings indicate that a broad range of predators actively or opportunistically exploit mouse lemurs, making them the most heavily preyed-upon lemur genera (Fichtel, 2016; Goodman and Ganzhorn, 2022).

**Table 1.**
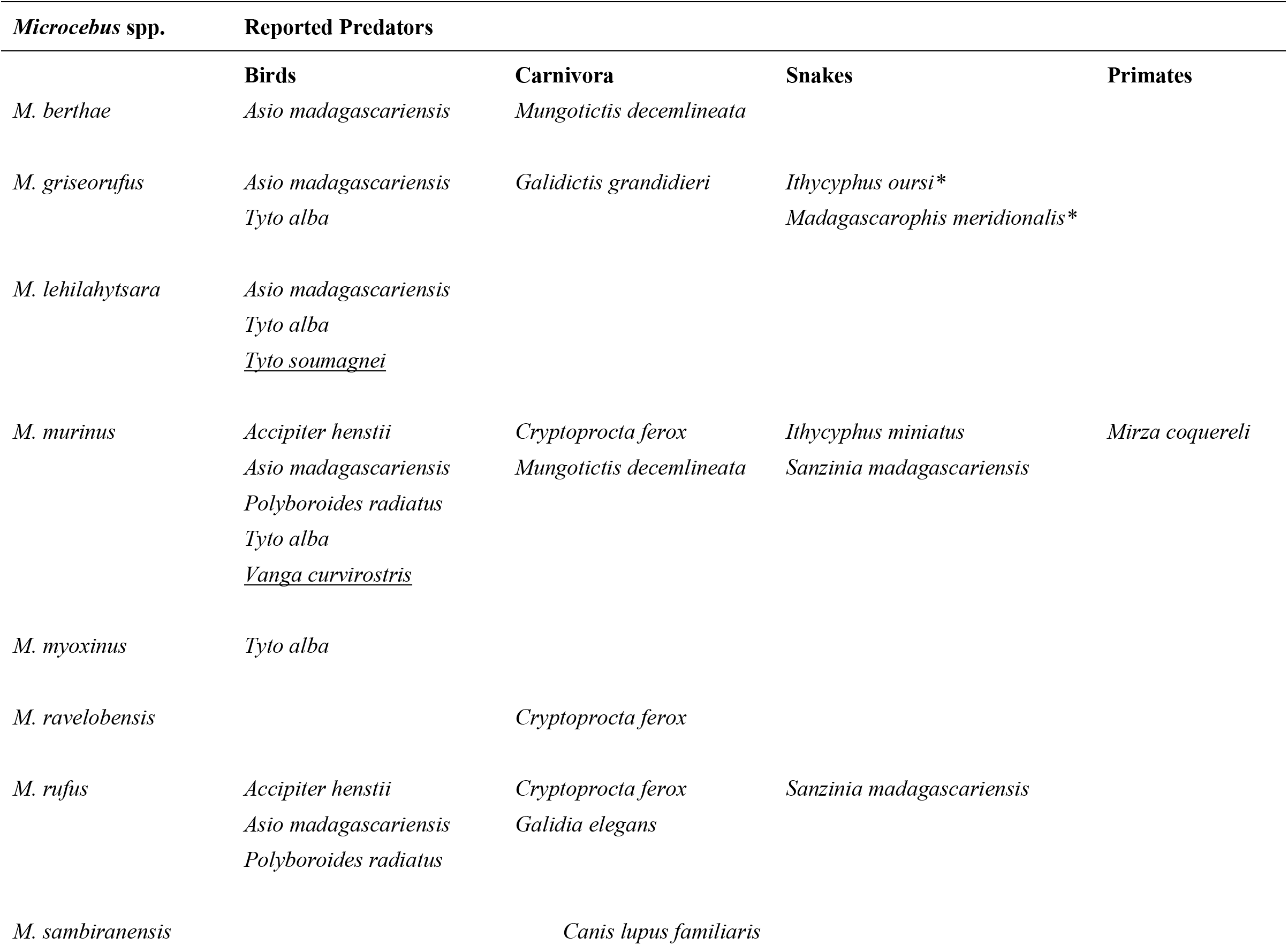
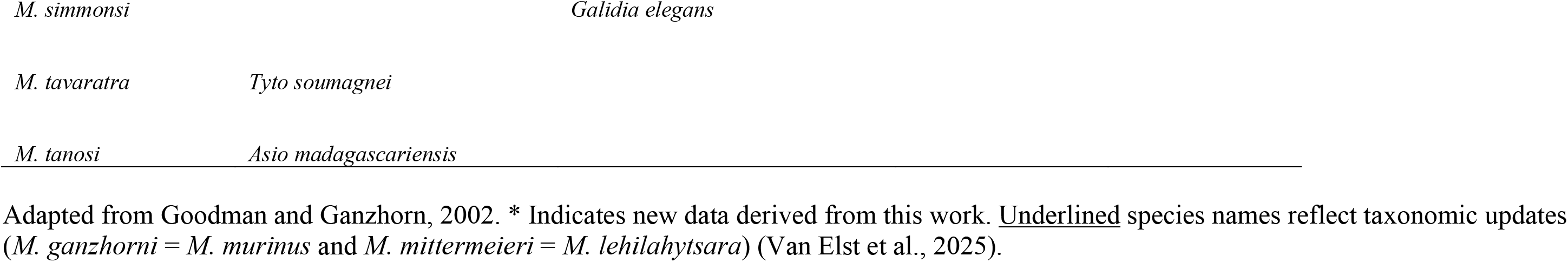
Summary of known *Microcebus* predators.

Whilst *Microcebus* spp. are strictly nocturnal, predators include both nocturnal and diurnal taxa. Based on the available literature (summarized in Goodman and Ganzhorn, 2022), the principal predators of *Microcebus* spp. comprise predatory raptors (e.g., *Accipiter henstii*, *Asio madagascariensis*, *Polyboroides radiatus* and *Tyto* spp.), Eupleridae (e.g., *Cryptoprocta ferox*, *Galidia elegans*, *Galidictis grandidieri*, and *Mungotictis decemlineata*), introduced carnivorans, and snakes (*Ithycyphus miniatus* and *Sanzinia madagascariensis*). Rare instances of predation by other primates and by humans have also been reported (Schliehe-Diecks et al., 2010; Borgerson et al., 2022). Since mouse lemurs occur in all forest types across Madagascar, spatial overlap with particular predator species varies among localities and taxa. Nevertheless, several documented predators (e.g., *Tyto alba*, *Cryptoprocta ferox*, and *Sanzinia madagascariensis*) have similarly broad geographic distributions, indicating that predation pressure on *Microcebus* is likely to be widespread.

Here, we present predation data on *Microcebus* spp. from two complementary sources in Andohahela National Park. First, direct evidence was obtained from a larger study of radio-collared mouse lemurs (n = 56) monitored at two sites between May–August and October–December 2024. Second, indirect evidence of predation was derived from the examination of pellets and prey remains collected at several owl roosts. Owl pellets provide a valuable record of small-vertebrate predation because they contain identifiable skeletal remains at fixed roost sites, enabling the detection of cryptic, nocturnal prey such as mouse lemurs even when predation events are rarely observed directly. Taken together, these data provide insight into predation pressure and the ecological dynamics influencing mouse lemurs in Andohahela.

The Andohahela protected area (76,020 ha), first designated as a Réserve Naturelle Intégrale in 1939 and renamed as a Parc National in 1998, is divided into three non-contiguous parcels spanning a steep climatic gradient (Goodman et al., 2018). Parcel I (62,093 ha), the largest, is dominated by lowland to medium altitude moist evergreen forest of the southern Anosyenne Mountains (Gautier et al., 2018). This mountain chain generates a dramatic rain shadow effect, reducing rainfall and humidity in the western portions of the park. Parcel II, by contrast, is smaller (13,574 ha) and with a distinctly xeric vegetation known as dry spiny thicket. A third and geographically separate Parcel III (437 ha), comprised primarily of low-elevation transitional forest, is situated further to the south (Figure 1). Mouse lemurs are found throughout the park, with *M. murinus* and *M. griseorufus* occurring in sympatry across the dry spiny thicket and transitional habitats (Parcels II and III), whereas *M. tanosi* occupies the moist evergreen forests (Parcel I) (Mittermeier et al., 2023).

**Figure 1.**
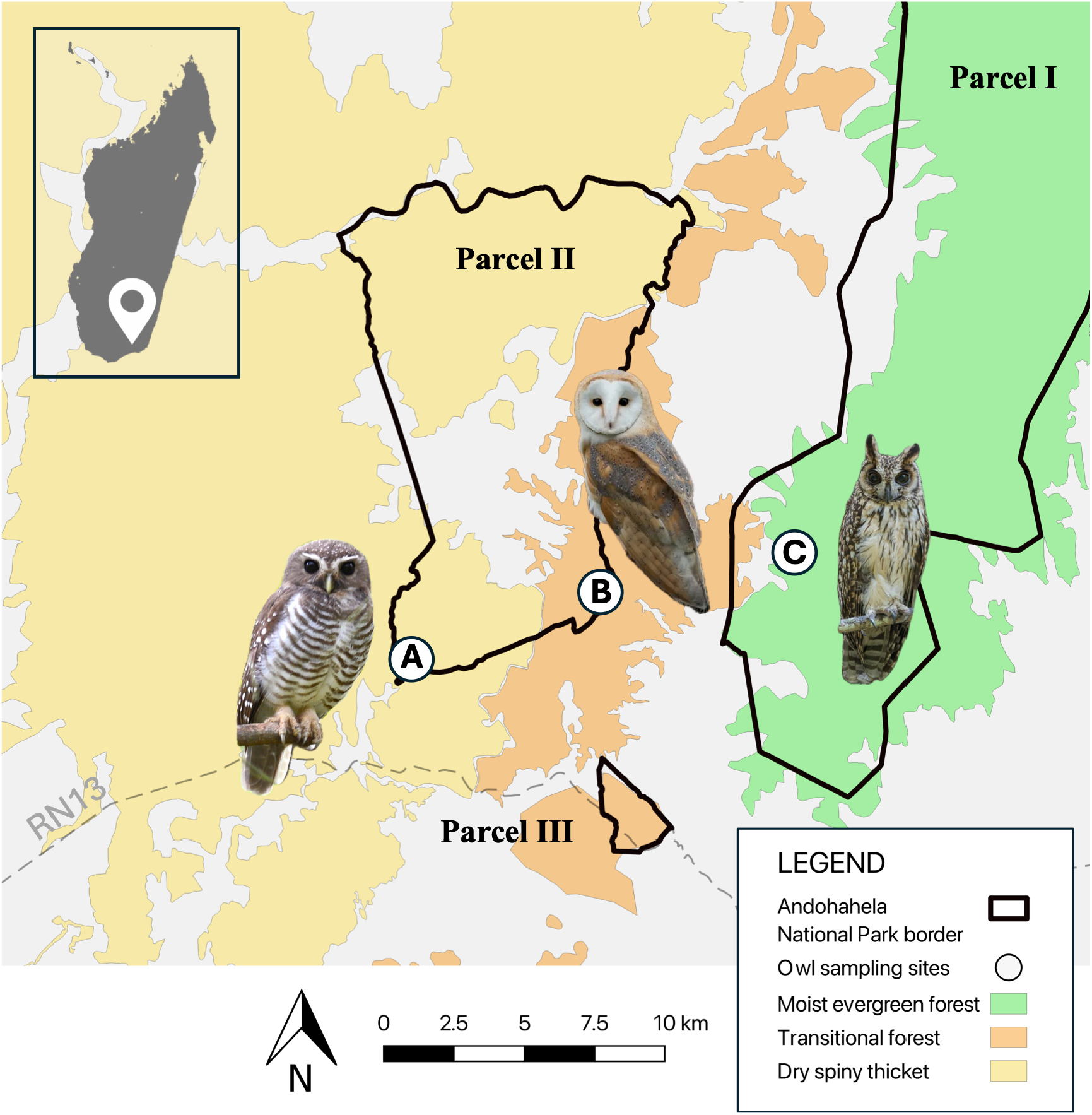
Owl sampling localities in Andohahela National Park. **A**) Mangatsiaka (*Athene superciliaris*). **B**) Tsimelahy (*Athene superciliaris* and *Tyto alba*) and **C**) Ebosika (*Asio madagascariensis*).

## Methods

### Study sites

#### Mangatsiaka

Mangatsiaka is a dry spiny thicket site located in the westernmost part of Parcel II (24° 58.064′ S; 46° 33.195′ E). The site receives relatively low annual rainfall (< 800 mm) and as a result, vegetation is dominated by dry spiny thicket, particularly species of Didiereaceae and Euphorbiaceae. Structurally, the forest is relatively homogeneous, although extensive cyclone damage is evident. Mean canopy height is approximately 6 m, with emergent *Alluaudia* spp. reaching up to 12 m. Narrow bands of gallery forest (e.g., *Tamarindus indica* and *Breonadia salicina*) occur along small river tributaries that traverse the site.

#### Tsimelahy

Tsimelahy is situated on the eastern edge of Parcel II, within the transitional forest zone (24° 56.705′ S; 46° 37.847′ E). Vegetation includes species characteristic of dry spiny thicket and transitional forest formations (e.g., *Commiphora brevicalyx*, *Dypsis decaryi*, and *Pachypodium lamerei*). The area is characterized by the sub-arid climate and receives an annual rainfall of approximately 830 mm, most of which falls during the wet season (November–April). The forest has a relatively open, homogenous structure with a low canopy height of 6–8 m, with emergent trees reaching up to 12 m (Goodman et al., 2018). The site is along the Tarantsy River, which originates in the Anosyenne Mountains to the east and drains westward into Lake Anony, providing an important refuge for a diverse range of animal species. The river margins are lined with relatively intact gallery forest.

#### Ebosika

Ebosika is a moist evergreen forest site located approximately 4 km east of the village of Tsimelahy, in the southwestern foothills of the Anosyenne Mountains (24° 56.639′ S; 46° 40.041′ E). The Ebosika forest is characteristic of Parcel I, with mean canopy height of 20–30 m and emergent trees reaching up to 30 m. Classified as lowland moist evergreen forest, the site supports a rich flora and fauna typical of south-eastern Malagasy evergreen moist forests (Goodman et al., 2018). Annual rainfall is approximately 1,355 mm. Although degraded at lower elevations, forest structure remains dense and relatively intact at higher altitudes and closer to the source of the Tarantsy River.

#### Telemetry-based detection of predation

Predation events on radio-collared mouse lemurs were documented during a concurrent telemetry study conducted at Mangatsiaka and Tsimelahy between May–August (dry season) and October–December 2024 (wet season). A total of 56 mouse lemurs were fitted with lightweight, thermistor-equipped VHF radio collars (ATS M1420; Advanced Telemetry Systems, Isanti, MN, USA; ~1.5 g). Collar deployment was balanced across species (*Microcebus murinus* and *M. griseorufus*) and sexes at both sites. Animals were monitored three to four times per week using standard radio-telemetry techniques (Yagi antennae and ATS receivers), with each individual followed for a minimum of 3–4 full-night tracking sessions. Species identities were confirmed from DNA extracted from xxx using the Qiagen DNeasy Blood & Tissue Kit. The mitochondrial gene cytochrome *b* was amplified and sequenced using primate primers L14724 and H15915 (Irwin et al., 1991).

#### Owl roosts and prey remains

At Mangatsiaka, pellet remains was collected from two *Athene superciliaris* roost sites approximately 70 m apart and used by the same pair of owls. Collections were made on 8 July 2024 and 20 November 2024. The primary roost site comprised a single perch at approximately 3 m above ground, beneath a densely tangled canopy. The second auxiliary site was located nearby in a small stream bed at a height of approximately 2.2 m above ground.

At the Tsimelahy site, assorted *Tyto alba* prey remains were collected on 9 July 2024 from an active roost site (170 m a.s.l.) located within 10 m of a small semi-permanent tributary of the Tarantsy River. Owl identity was confirmed by direct observation and the presence of dropped feathers. The roost was concealed approximately 10 m above ground in a *Dypsis decaryi* palm, with pellet remains scattered below among *Agave sisalana* plants. Additional prey remains were recorded at an abandoned roost site approximately 60 m to the south that had been damaged by Cyclone Batsirai in 2022; material remaining at this site was identified from photographs (Figure 2E). Further material was collected on 30 June 2024 from an active *Athene superciliaris* roost within transitional forest. This roost, occupied by two adult owls, was situated beneath the canopy at approximately 3 m height in a *Commiphora* tree.

**Figure 2.**
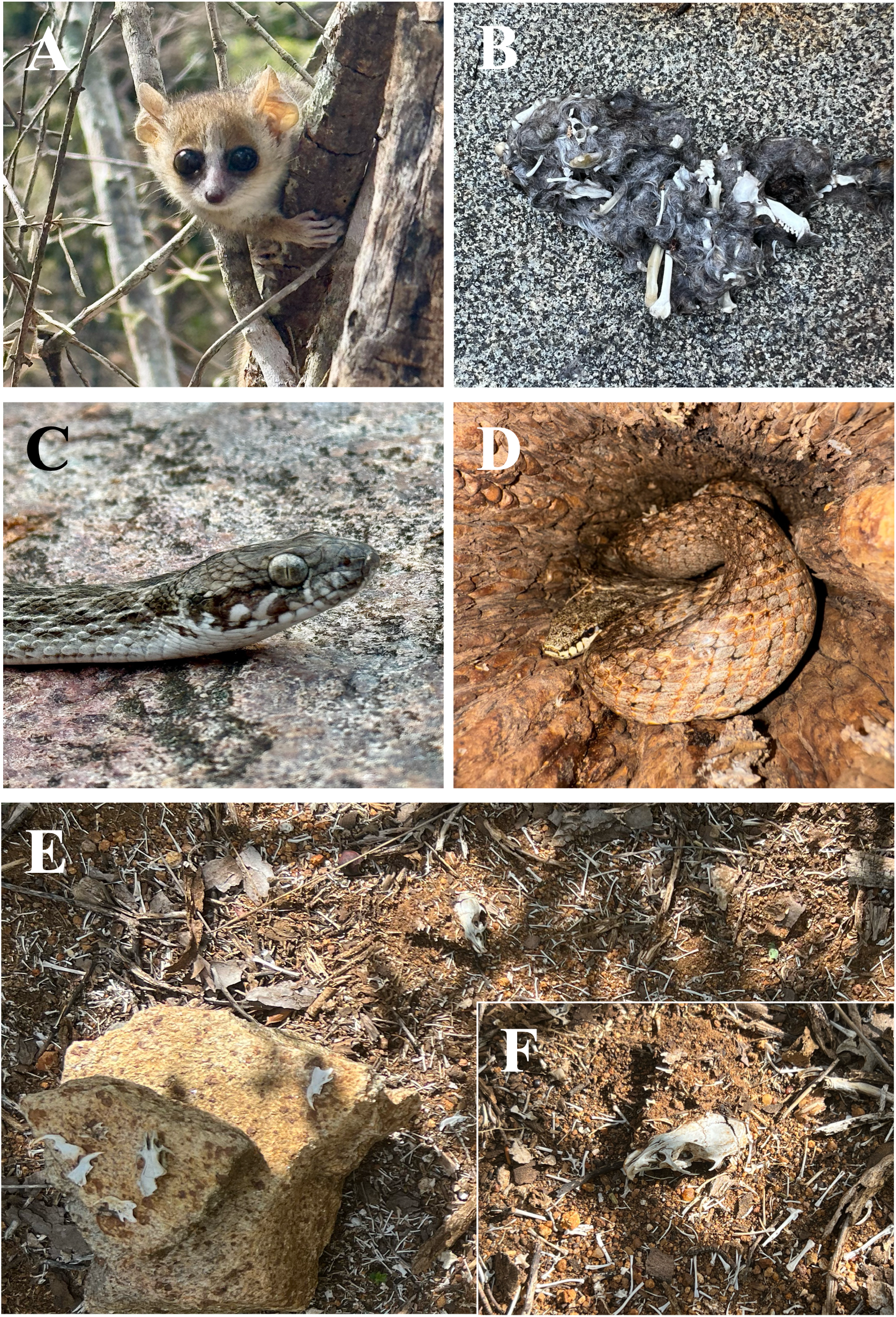
**A**) *Microcebus griseorufus*, **B**), *Asio madagascariensis* pellet containing regurgitated *Microcebus* remains at Ebosika. **C**) *Madagascarophis meridionalis*, **D**) *Ithycyphus oursi*, **E**) *Tyto alba* roost debris site, Tsimelahy. **F**) *Rattus rattus* crania from site in E.

In the humid forest at Ebosika, two pellets attributed to *Asio madagascariensis* were collected. The first pellet, consisting of a complete skeletonised small mammal, was recovered on 11 June 2023 from a large, exposed boulder (750 m a.s.l.) in the centre of a partially dry riverbed (Figure 1B). No additional remains were found in the immediate vicinity; however, an adult *A. madagascariensis* was observed roosting within approximately 150 m of the deposit. A second pellet containing a single small mammal was located and collected on 1 December 2024 approximately 200 m away in leaf litter beneath a large *Canarium madagascariensis* tree. The size and composition of these pellets are characteristic of *A. madagascariensis* and do not correspond to pellets produced by other owl species known from the area.

#### Bone identification

Field sample collection and preparation largely followed Yalden (2020). Owl pellets were separated and soaked in soapy water for 24 h to soften and facilitate disassembly, then rinsed through a fine-mesh sieve. Sorted elements were transferred to aluminium trays containing a small amount of water and, using fine tweezers, skeletal and dental remains were carefully removed and placed on absorbent paper for drying. Identification of skeletal remains was based primarily on morphological criteria and conducted by authors V.J.N.H. and S.M.G. When genus or species-level attribution was not possible, remains were classified at the lowest reliable taxonomic level. Minimum number of individuals (MNI) was determined using paired and diagnostic skeletal elements; for amphibians, clavicles were used as reference elements, whereas for mammals MNI was estimated from the number of skulls and/or mandibles.

## Results

### Predatory observations

At Mangatsiaka, we confirmed the predation of two adult mouse lemurs by snakes. One individual was an adult female *Microcebus griseorufus* and the second was an adult female *M. murinus*. Species identity of both individuals was confirmed genetically, and both had been fitted with thermistor-equipped VHF radio collars. The *M. griseorufus* (52 g when captured) was tracked to an underground termite nest, where its signal remained stationary for several days before the site was excavated on 5 May 2024. The individual had been consumed by *Madagascarophis meridionalis* (total length [TL] = 750 mm; mass = 211 g), which was located approximately 50 cm below ground in a small cavity formed by tree roots. The *Microcebus murinus* (89 g when captured) was recovered from within the lower trunk of a large *Opuntia stricta* cactus located along the margin of a dry streambed. After several days of signal inactivity and abnormally low recorded body temperature, inspection revealed that the individual had been predated by *Ithycyphus oursi* (TL = 1361 mm; mass = 455 g). In both cases, predation was confirmed by the presence of the radio collars within the digestive tracts of the snakes. Tags were later recovered from snake faeces. These observations provide direct evidence of snake predation on adult mouse lemurs in the dry spiny thicket of Mangatsiaka. Snake identification was based on morphological characters and field familiarity with the species.

Owl pellets, prey remains and regurgitated material were collected from three sites and associated with three owl species: *Asio madagascariensis* at Ebosika, *Tyto alba* at Tsimelahy, and *Athene superciliaris* at Mangatsiaka and Tsimelahy (Table 2). At Ebosika, two pellets attributed to *Asio madagascariensis* contained the remains of a single mouse lemur (*Microcebus* sp.) and the other of a single black rat (*Rattus rattus*). Given the locality and habitat type (deep within moist evergreen forest of Parcel I), the predated mouse lemur was inferred to be *M. tanosi*. Material recovered from *T. alba* roosts at Tsimelahy consisted exclusively of small mammals and amphibians (Table 2). Mammalian prey comprised two rodent species (*R. rattus* and *Mus musculus*) and one shrew (*Suncus murinus*), all three genera introduced to Madagascar, and together accounted for 27.4% of the MNI and 76.3% of total prey biomass. Amphibian remains were tentatively attributed to *Boophis doulioti*, *Laliostoma labrosum*, and *Ptychadena mascareniensis* based on osteological size and known local occurrence. Collectively, amphibians comprised 23.7% of prey biomass, with *P. mascareniensis* representing the most frequently recovered amphibian taxon.

**Table 2.**
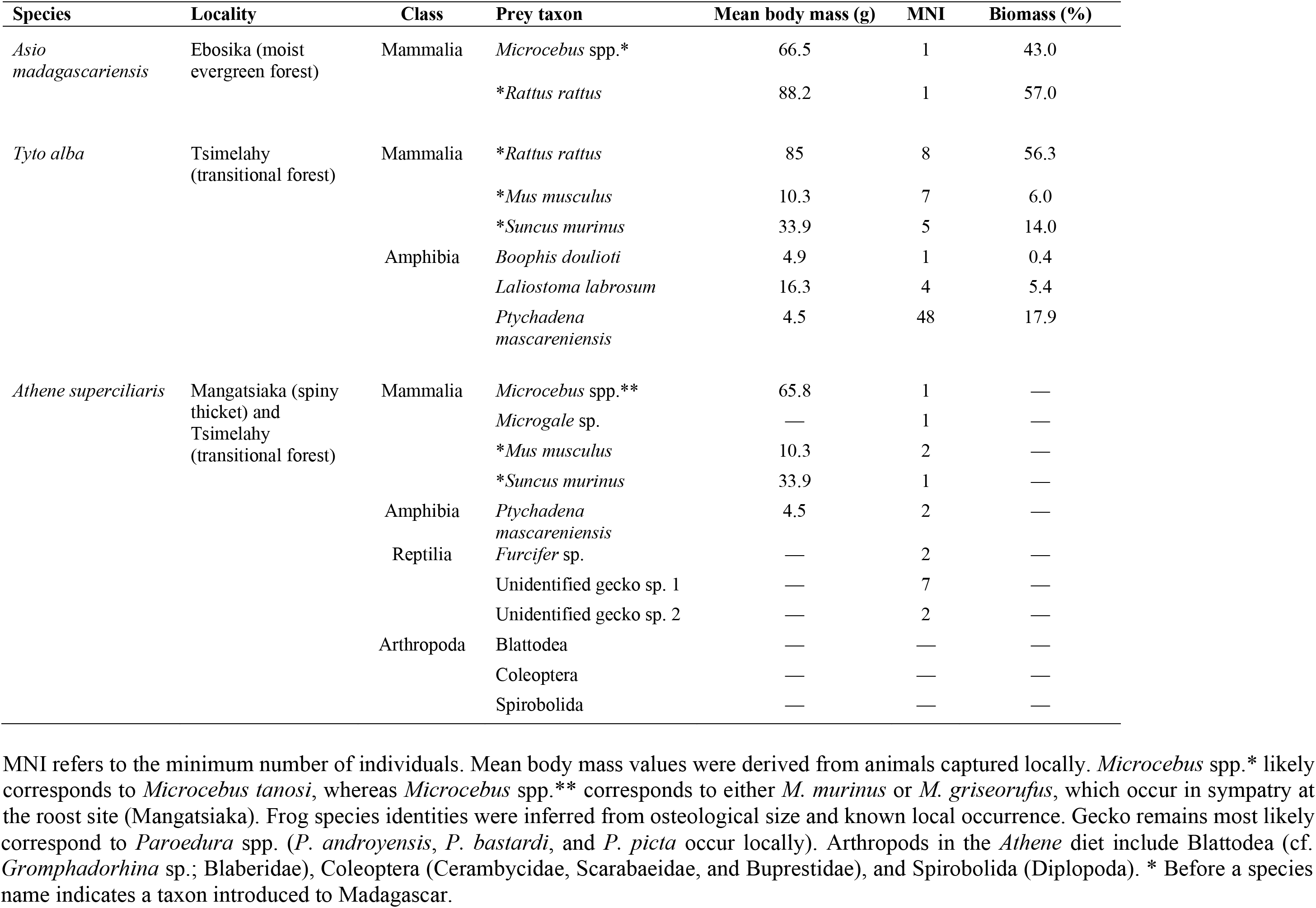
Dietary composition of three owl species in Andohahela National Park, south-eastern Madagascar.

In contrast, material recovered from *Athene superciliaris* roosts at Mangatsiaka and Tsimelahy (pooled) was composed predominantly of invertebrate remains, for which minimum numbers of individuals could not be reliably estimated due to high fragmentation of the recovered body parts (Table 2). Mammalian prey included *Microcebus* sp., *Microgale* sp., and *Suncus murinus*, each represented by a single individual, and *Mus musculus*, represented by two individuals. Amphibian remains were limited to two individuals, most likely attributable to *Ptychadena mascareniensis*. Reptile remains included a single chameleon (*Furcifer*) and two gekkonid taxa (provisionally assigned to *Paroedura* based on local abundance). Arthropod prey included Blattodea (cf. *Gromphadorhina* sp.; Blaberidae), Coleoptera (Cerambycidae, Scarabaeidae, and Buprestidae), and Spirobolida (Diplopoda). No estimates of biomass were calculated for invertebrate prey.

## Discussion

This study documents two additional endemic snake species as predators of *Microcebus*, *Ithycyphus oursi* and *Madagascarophis meridionalis*. *Ithycyphus oursi* is considered widespread in southern and southwestern Madagascar (Glaw and Vences, 2007) and is ecologically and behaviourally similar to its congener *I. miniatus*, a recognised mouse lemur predator from the north-western region of the island (Richard, 1978). *Ithycyphus oursi* is a large (total length up to 1568 mm), aggressive and mildly venomous predator with a broad range of prey types. The species regularly occupies tree holes and likely constitutes a significant predator of mouse lemurs across its range. In contrast, predation by *Madagascarophis meridionalis* is somewhat unexpected given its largely terrestrial habits and smaller body size (total length up to 870 mm). Although the species is among the most common snakes in southern Madagascar (Vences, 2011), its diet predominantly comprises small vertebrates such as amphibians and lizards. Nonetheless, occasional predation on larger vertebrate prey has been reported (Neaves et al., 2019), indicating greater dietary flexibility than previously assumed. It cannot be ruled out that this predation event involved scavenging a dead *Microcebus*; however, during the telemetry work, this snake species was observed high in a tree, unsuccessfully striking at a passing mouse lemur. Similar failed attempts by its congener *M. colubrinus* to predate young *Cheirogaleus medius* are also documented (Dausmann, 2010).

In contrast to avian predators, whose diets can be assessed through accumulated remains at roost sites, predation by snakes is typically documented only through opportunistic observations. In the present study, snake predation events were detected solely because prey individuals were fitted with VHF radio collars. It is therefore likely that snake predation on mouse lemurs is substantially underestimated. Density data are lacking for most Malagasy snake species, although both *Ithycyphus* and *Madagascarophis* are considered relatively common across much of their ranges.

In contrast to endothermic predators such as *Cryptoprocta ferox* and *Tyto alba*, which require frequent meals, a single ~60 g prey item may sustain a medium-sized snake such as *I. oursi* for an extended period. In our study, the interval between the disappearance of a radio-collared mouse lemur and the subsequent defecation event by the predatory snake suggests a digestion period of approximately nine days. Although based on a single observation, this provides a rare field-based estimate of digestion time following consumption of a small primate and underscores the difficulty of inferring predation rates from snake encounter frequencies alone. Furthermore, whereas owls are largely restricted to predating mouse lemurs during their nocturnal activity phase, snakes such as *I. oursi* and *I. miniatus* are likely to encounter mouse lemurs not only while they are active but also at diurnal sleeping sites and during periods of torpor. Consequently, mouse lemurs may experience predation risk throughout the daily and annual cycles, including during periods of reduced activity, highlighting snakes as potentially important but under-recognized predators of small primates.

In addition to predation by snakes, avian predators also represent an important component of the predator assemblage affecting mouse lemurs in Andohahela. This study confirms *Asio madagascariensis* as a predator of mouse lemurs in southern humid forest habitats (Goodman and Ganzhorn, 2022). At Ebosika, the *Microcebus* remains attributed to *A. madagascariensis* most likely correspond to *M. tanosi*, which occupies the moist evergreen forests of Parcel I, although *M. murinus* is also known to occur at lower elevations and forest edges (S. Hyde Roberts, unpubl. data). Previous work at the nearby Nahampoana Reserve reported that 4.4% of *A. madagascariensis* dietary biomass (7.2% of prey individuals) consisted of *M. tanosi* (Goodman et al., 1993b), supporting the view that mouse lemurs can form a regular component of its diet in moist forest ecosystems. A second pellet associated with *A. madagascariensis* demonstrates the diet also contains invasive rodents (*Rattus rattus*), which are abundant in disturbed and degraded areas (Ganzhorn, 2003; Lehtonen, 2013).

The prevailing assumption that *Tyto alba* constitutes a principal predator of *Microcebus* species across Madagascar is well established and is intuitively supported by their shared nocturnal activity patterns and frequent habitat overlap. In the dry forests of Bezà-Mahafaly in the southwest, *M. griseorufus* represented 5.7% of total prey individuals and 22.9% of prey biomass in barn owl diets (Goodman et al., 1993a). However, our findings from Andohahela National Park do not support this pattern. Despite high local densities of mouse lemurs at Tsimelahy, we found no evidence that *Microcebus* formed part of the diet of *T. alba* at this locality. Although our material was derived from only two roost sites, no mouse lemur remains were detected during two sampling events conducted several months apart. At Bezà-Mahafaly, seasonal variation in predation rates on *Microcebus* has been reported (Goodman et al., 1993a); however, because our sampling encompassed multiple months, it is unlikely that mouse lemur predation was simply missed due to seasonal effects.

Instead, *T. alba* diets at Tsimelahy were dominated by rodents and amphibians, suggesting a strong local prey preference. Notably, both roost sites were located on the margins of a small tributary of the Tarantsy River, and the owl was frequently observed hunting along this riparian corridor. Such prey may be more vulnerable to capture because they occupy more exposed microhabitats (e.g., open stream margins and ephemeral pools), whereas mouse lemurs may exploit structurally complex vegetation such as spiny thicket and adopt effective antipredator strategies (Anderson, 1998; Fichtel, 2016). Notably, these results are consistent with findings from the nearby littoral forest of Petriky, where mouse lemurs likewise comprised only a small proportion of barn owl diets (Goodman et al., 2025), and from the eastern rainforest site of Andasibe (Goodman et al., 1993a). It has further been suggested that in areas where *Rattus rattus* dominates the small mammal community, *A. madagascariensis* is more likely to prey upon rodents than on *Microcebus* (Langrand and Goodman, 1996). Our results from Tsimelahy are consistent with this hypothesis and indicate that barn owl predation on mouse lemurs may be highly context-dependent, varying with prey availability and habitat structure.

In contrast to both *A. madagascariensis* and *T. alba*, dietary evidence from *Athene superciliaris* indicates a markedly different predatory strategy. Despite its wide distribution in southern Madagascar, the diet of *A. superciliaris* remains poorly known. Material recovered from *A. superciliaris* roosts in this study was dominated by invertebrate remains, with vertebrate prey occurring only rarely and comprising a taxonomically diverse assemblage of small mammals, amphibians, and reptiles. Partial remains of a single mouse lemur (*Microcebus* sp.) were identified among the collected mammalian materials, representing the first documented record of mouse lemur predation by *A. superciliaris*. However, its low representation relative to other prey items suggests that this species is unlikely to constitute a regular or specialized predator of mouse lemurs. Instead, *A. superciliaris* appears to be an opportunistic generalist, exploiting locally abundant and easily captured prey such as invertebrates (Blattodea, Coleoptera and Spirobolida), small reptiles, and amphibians. The predominance of invertebrates, combined with the small body size and hunting mode (perch hunting, sit-and-wait strategy) of *A. superciliaris*, further supports the interpretation that predation on mouse lemurs is not particularly common. Camera traps deployed at the roost site in Tsimelahy also documented simultaneous use of the site by both the owl and mouse lemurs at close proximity, without evidence of antagonistic interactions. Together, these findings suggest that smaller owls may contribute to background predation pressure on *Microcebus* populations but are unlikely to represent a major selective force compared with larger avian or reptilian predators.

While there is no doubt that raptors and carnivores exert substantial population-level predation pressure on mouse lemurs (Goodman et al., 1993a, 1993b; Goodman and Ganzhorn, 2022), we postulate that snakes also represent a currently underappreciated component of the predator guild. At Bezà-Mahafaly, an estimated 500 mouse lemurs are consumed annually by 9–10 pairs of *T. alba*, and up to 4.5% of the mouse lemur population may be lost to diurnal raptors in the moist evergreen forests of Ranomafana (Karpanty, 2003). In the dry deciduous forest of Kirindy (CNFEREF), *Cryptoprocta ferox* is estimated to prey upon approximately 18% of the *Microcebus* population each year (Hawkins and Racey, 2008). In contrast, no comparable estimates exist for predatory snake species. Nevertheless, our observations suggest that snakes contribute to mouse lemur mortality. Indirect support for this hypothesis is provided by the pronounced antipredator behaviours directed toward snakes observed in *Microcebus* and other lemurs (Schülke, 2001; Deppe, 2006; Eberle and Kappeler, 2008; Rahlfs and Fichtel, 2010), indicating strong selective pressure from reptilian predators. Such pressure is unlikely to be restricted to the smallest lemurs alone (Goodman and Ganzhorn, 2022).

Whilst our study documents snake and owl predation on mouse lemurs, several additional cheirogaleid predators are known to occur within Andohahela. *Cryptoprocta ferox* has previously been reported from Mangatsiaka and is believed to have been abundant in past decades; however, no individuals were detected during the present study, including in camera traps image, and local knowledge suggests that the population has declined in recent years (N. J. Ralatiany, pers. comm., 2024). Predatory birds such as *Polyboroides radiatus* and *Vanga curvirostris* were confirmed at both Mangatsiaka and Tsimelahy, while *Tyto soumagnei*, *Sanzinia madagascariensis*, and *Galidia elegans* are known to occur in the moist evergreen forest at Ebosika. A notably smaller 4th owl species, *Otus rutilus*, was present at all sites but is not known to prey on mouse lemurs. Together, these observations indicate that the local predator community is diverse and that the predation events documented here likely represent only a subset of the predator–prey interactions affecting *Microcebus* populations. This underscores the difficulty of quantifying predation pressure in small, nocturnal primates and highlights the importance of integrating multiple lines of evidence when assessing predator impacts.

## Acknowledgments

We are grateful to the Malagasy authorities, including Direction des Aires Protégées des Ressources Naturelles Renouvelables et des Ecosystèmes, which kindly granted the necessary research authorizations from which the presented herein originate. We thank Madagascar National Parks for facilitating our research and our wider Malagasy team who helped make this work possible. This research was supported by NSFDEB-2148914 to ADY. Finally, we thank the IUCN SOS Lemur Fund for supporting the work of VJNH. Misaotra betsaka.

## Ethics Statement

All field procedures involving animals were conducted in accordance with relevant national and international guidelines for the ethical treatment of wildlife. Field research was conducted under permits granted by the Malagasy Ministry of Environment and Sustainable Development through the Direction des Aires Protégées des Ressources Naturelles Renouvelables et des Ecosystèmes (167/23/MEDD/SG/DGGE/DAPARNE/SCBE.Re, issued 12 May 2023; and permits. 096/24 and 341/24/MEDD/SG/DGGE/DAPARNE/SCBE.Re, issued 27 March and 13 September 2024). All animal handling and sampling procedures were approved by the Duke University Institutional Animal Care and Use Committee (IACUC; protocol no. A163-22-09) and were designed to minimize stress and disturbance. The use of radio collars and tissue sampling was justified by the need to investigate ecological and behavioural processes, and all individuals were released at the site of capture following processing.

## Conflict of Interest

The authors declare no conflict of interest.

## Author Contributions

A.D.Y. and S.H.R. conceived the broader study. S.H.R., J.C.S., and A.D.Y. designed the research. Field data were collected by S.H.R., J.C.S., V.J.N.H., and E.R. Prey identifications were conducted by V.J.N.H., S.M.G., and B.R. Data analysis was performed by S.H.R. The manuscript was written by S.H.R. with input from all authors. All authors approved the final version of the manuscript.

## Data Availability

All data generated or analysed during this study are included in this article. Further enquiries can be directed to the corresponding author.

## Notes

### Competing Interest Statement

The authors have declared no competing interest.

